# Pathway centered analysis to guide clinical decision-making in precision medicine

**DOI:** 10.1101/2021.04.30.442131

**Authors:** Luís B. Carvalho, J. L. Capelo, Carlos Lodeiro, Rajiv Dhir, Luis Campos Pinheiro, Mariana Medeiros, Hugo M. Santos

## Abstract

Changes in the human proteome caused by disease before, during and after medical care is phenotype-dependent, so the proteome of each individual at any time point is a snapshot of the body’s response to disease and to disease treatment. Here, we introduce a new concept named differential Personal Pathway index (dPPi). This tool extracts and summates comprehensive disease-specific information contained within an individual’s proteome as a holistic way to follow the response to disease and medical care over time. We demonstrate the principle of the dPPi algorithm on proteins found in urine from patients suffering from neoplasia of the bladder. The relevance of the dPPi results to the individual clinical cases is described. The dPPi concept can be extended to other malignant and non-malignant diseases, and to other types of biopsies, such as plasma, serum or saliva. We envision the dPPi as a tool for clinical decision-making in precision medicine.

## Main

The advent of high-resolution mass spectrometry-based proteomics in conjunction with advanced bioinformatics tools has made it possible to quantify thousands of proteins in a straightforward, reproducible and robust way, with less and less recourse to time-consuming methods involving labelling, protein standards or calibration curves^1^. This remarkable progress has made it possible to follow the biochemical changes at the protein level in a given individual from the onset of disease^2^.

To express the variation in an individual’s proteome over time in a way that is amenable to personalized medicine, we propose using the dPPi, calculated from the numerical output of proteomics analysis of any type of medical biopsy at different time points, as follows:

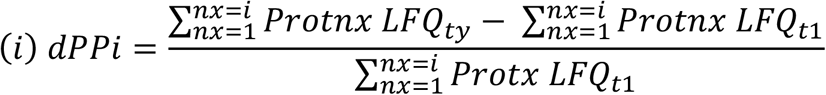

where 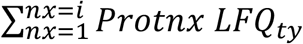 accounts for the sum of all label-free quantitation (LFQ) values of proteins of interest at a given time *t_y_* whilst 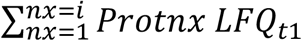 accounts for all LFQ measurement of the same proteins at the time of disease onset *t*_1_.

Based on variation in the levels of tens of signature proteins, the dPPi will reflect both the response of the individual’s proteome to whatever course the disease takes and any medical care that ensues. The variation is expected to reflect the status of the biochemical pathways to which the proteins correspond, specifically chosen to combine multiple sources of diagnostic information. Thus, evolution of the disease is monitored in a comprehensive yet simple manner. Essentially, the more positive the dPPi, the worse the course of disease; the more negative the dPPi, the better the clinical outlook. The dPPi can therefore flag the need for intervention when a disease is progressing. This concept is demonstrated below with three examples of urinary proteomes from patients with bladder cancer (BC).

First, we selected a group of BC patients with multiple recurrences, progression to T2-stage BC, and in some cases, patients who died due to BC (Group A, Advanced). The second group consists of early T1-stage cancer patients with no record of medical intervention at the time of diagnosis or when the first urine was taken (Group I, Initial). The clinical information about both groups is presented in Table 1 supplementary material (Table SM1).

Next, we cross-checked the proteins that were differentially expressed between the two groups (n = 71, Fig. 1A, 1B and 1C and Table SM 2) against the Hanahan and Weinberg biological hallmarks of cancer (HWhc) and the corresponding biochemical pathways (Fig. 1E and 1F^5,6^). This allowed us to investigate how the status of these pathways changed over time by following some patients with BC for up to 40 months.

**Figure 1.**
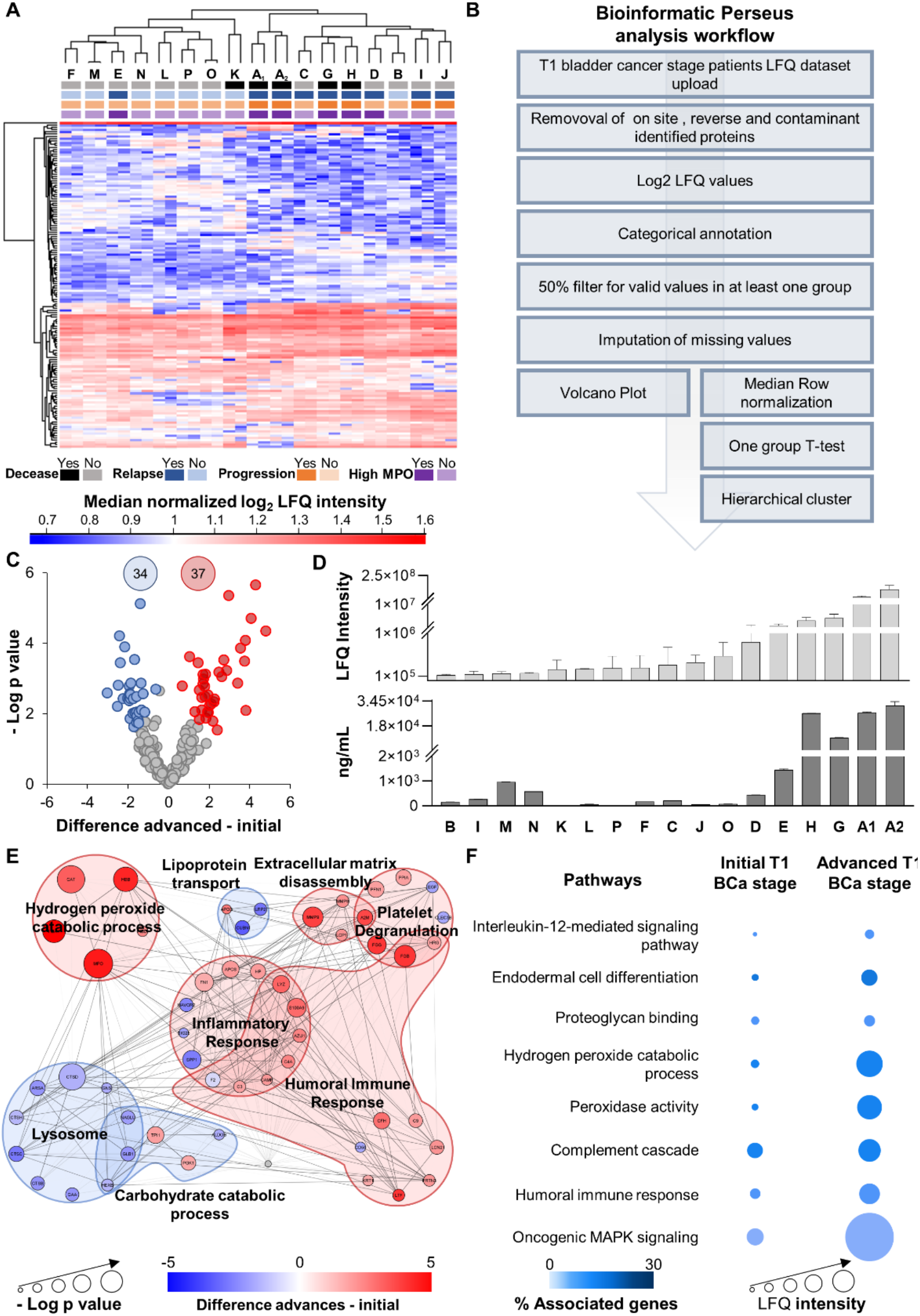
**a**, Hierarchical clustering of T1-BC patients into two groups, as expected, initial and advanced (average linkage, no constraint, pre-processing with k-means, and Euclidean distance between column trees). **b**, Perseus Workflow used to obtain the hierarchical clustering and the volcano plots (Perseus v. 10.6.10.5). **c**, Volcano plots showing statistically significant changes in protein levels in the A group (high frequency relapses and progression) compared to the I group (Initial stage) of T1-BC patients, according to the Student’s t-test (FDR 0.05 and S_0_ of 0.1). Dots represent proteins showing statistically significant increases (red), decreases (blue) or non-statistically significant changes (gray). Total numbers of proteins showing statistically significant differential expression are shown in the corresponding-colored circles. **d**, Validation of mass spectrometry data using myeloperoxidase (MPO) protein label free quantification (LFQ) and ELISA (ng/mL). MPO protein has been proposed recently as a marker of poor prognosis in lung^3^ and ovarian^4^ carcinomas. To the best of our knowledge there are no studies reporting overexpression of MPO as a prognosis tool for BC carcinoma. **e**, Protein-protein interaction analysis of the proteins showing statistically significant differential expression as derived from the volcano plot in **c** (proteins are listed in Table 1SM). **f**, Pathways selected for calculating the Personal Pathway Index, after applying the guidelines provided by Hanahan and Weinberg^5,6^ to the proteins showing statistically significant differential expression as derived from the volcano plot in **c** (Table 2SM). Differentially expressed proteins were grouped according to their functions and networks were analyzed in ClueGo, Cytoscape (v. 3.8.0. 8). The LFQ average for all proteins involved in the pathway is expressed as a circle area plot.

Patient I presented symptoms matching BC, and so one reference urine sample (B1) was taken before urinary and bladder exploration (Fig. 2). This patient was diagnosed with T1-Stage BC and TURBT was performed. One month later, a second analysis of the urinary proteome (B2) revealed a small difference of −4% in the dPPi since BC onset (Fig. 2E, B2 - B1), suggesting that the disease was progressing. Fig. 2E, B2 - B1, clearly shows that pathways have the same level of dysregulation than when the patient was operated. This prompted us to call the patient for follow-up. After inspection of the bladder the diagnosis was cystitis, so no surgical intervention was needed.

**Figure 2.**
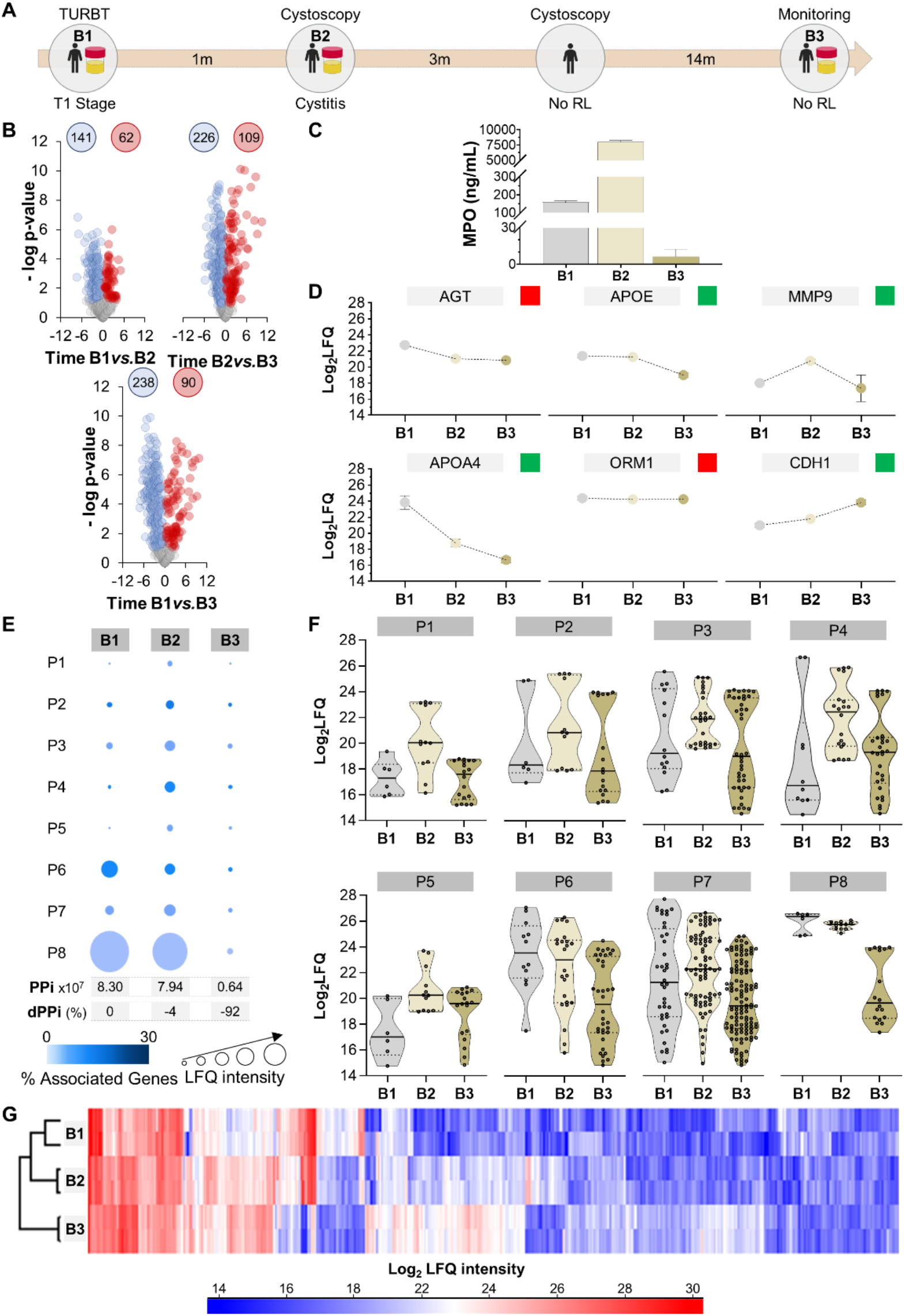
**a**, Timeline of disease course in Patient I shows intervals in months (m) between medical interventions and urine sampling (B1 to B3). TURBT: trans urethral resection of bladder tumor; Re-TURBT: repeated trans urethral resection of bladder tumor; RL: relapse. **b**, Volcano plots showing proteome changes in the urines sampled at times B1, B2, B3. Dots represent proteins showing statistically significant increase (red), decreases (blue) or non-statistically significant changes (gray) according to Student’s *t*-test (FDR 0.05 and S0 of 0.1). **c**, ELISA quantification of Myeloperoxidase (MPO, ng/mL) in urine samples. Myeloperoxidase (MPO, ng/mL) in urine samples. **d**, Variation for six known protein biomarkers for bladder cancer: Angiotensinogen (AGT)^7^, Apolipoprotein E (APOE)^8^, Matrix metalloproteinase-9 (MMP9)^9^, Apolipoprotein A-IV (APOA4)^10^, Alpha-1-acid glycoprotein 1 (ORM1)^11^ and Cadherin-1 (CDH1)^12,13^. Green and red squares indicate whether the variation in the urine of Patient I accords or does not accord, respectively, with trends reported for each marker in the literature. **e**, Personal pathway index (PPi) at each time point and estimated differential PPI (dPPi) calculated as explained in the text. P1: Interleukin-12-mediated signaling pathway; P2: endodermal cell differentiation; P3, proteoglycan binding; P4: hydrogen peroxide catabolic process; P5: peroxidase activity; P6: Complement cascade; P7: humoral immune response; P8: Oncogenic MAPK signaling. **f**, Variation in protein LFQ values (including replicates) at each sampling point for each pathway assessed. **G**, Hierarchical clustering of the three urinary proteomes of Patient II. Protein LFQ values were used to perform the cluster analysis (with average linkage, no constraint, preprocessing with k-means and Euclidean distance between column tree).

This result again shows the utility of the dPPi as a tool to follow the personal variations in the urine proteome caused by disease. Another cystoscopy three months later revealed no evidence of relapse. Fourteen months later Patient II was monitored, and the urine taken at that time (Fig. 2E, B3) had a dPPi of −92% relative to BC onset, indicating clear down-regulation of the key immunological and inflammatory responses (Fig. 2E) and reflecting the good progression of this patient.

The second example is Patient II, an interesting case because of frequent relapses (Fig. 3). The first analysis of the urine proteome, the reference one, was done 64 months after onset of BC just before a TURBT was performed because of a relapse. Two months later a new urine analysis, revealed a +104%, change in the dPPi, with some of the pathways doubling their expression levels (Fig. 3E, F1 - F2). The increase in expression of the MAPK pathway proteins was particularly marked (P8, Fig. 3E). Based on this data, the patient was called for a urological assessment which revealed another relapse of the tumor, confirming the reason for the rise in the dPPi. Another TURBT was performed, and BC was documented.

**Figure 3.**
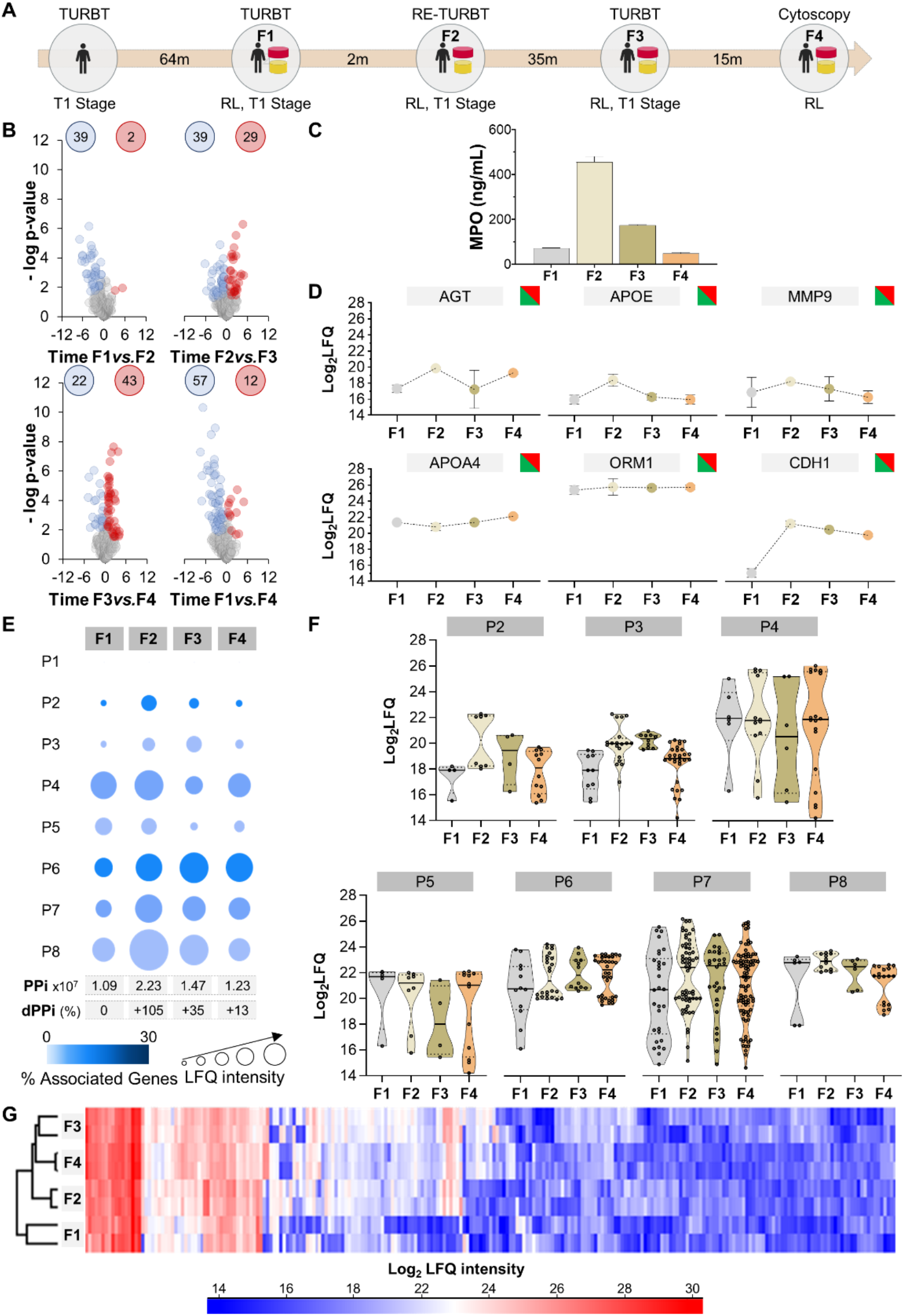
**a**, Timeline of disease course in Patient II (patient F) showing intervals in months (m) between medical interventions and urine sampling (F1-4). TURBT: trans urethral resection of bladder tumor; Re-TURBT: repeated trans urethral resection of bladder tumor; RL: relapse. **b**, Volcano plots showing proteome changes in urines collected at timepoints F1-4 as in **a**. Dots represent proteins showing statistically significant increases (red), decreases (blue) or non-statistically significant changes (gray) according to Student’s t-test (FDR 0.05 and S0 of 0.1). **c**, ELISA quantification of Myeloperoxidase (MPO, ng/mL) in urine samples. **d**, Variation in known protein biomarkers for bladder cancer Angiotensinogen (AGT)^7^, Apolipoprotein E (APOE)^8^, Matrix metalloproteinase-9 (MMP9)^9^, Apolipoprotein A-IV (APOA4)^10^, Alpha-1-acid glycoprotein 1 (ORM1)^11^ and Cadherin-1 (CDH1) ^12,13^. Green and red in squares indicate whether the variation in the urine of Patient II accords or does not accord, respectively, with trends reported for each marker in the literature. **e**, Personal pathway index (PPi) at each time point and estimated differential PPi (dPPi) calculated as explained in the text. P1: Interleukin-12-mediated signaling pathway; P2: endodermal cell differentiation; P3, proteoglycan binding; P4: hydrogen peroxide catabolic process; P5: peroxidase activity; P6: Complement cascade; P7: humoral immune response; P8: Oncogenic MAPK signaling. **f**, Variation in protein LFQ values (including replicates) at each sampling point for each pathway assessed. **G**, Hierarchical clustering of the four urinary proteomes for Patient II. Protein LFQ values were used to perform the cluster analysis (with average linkage, no constraint, preprocessing with k-means and Euclidean distance between column tree).

Subsequently, 35 months later (Fig. 3E, F3), during a routine surveillance assessment the urine proteome analysis revealed a dPPi of +35% relative to the original reference value, with increased levels of expression noted in most of the hallmark pathways. The patient was thus called again for a cystoscopy evaluation, which in line with the dPPi prediction showed relapse of BC. Consequently, a repeat TURBT was performed. Finally, a fourth urine proteomic analysis was performed 15 months later (Fig 3E, F4), which showed a change in the dPPi value of +14%, suggesting the tumor had relapsed again. This was later confirmed by cystoscopy and currently the patient is waiting for surgical intervention.

As a final example, patient III (Fig. 4) presented with symptoms of BC. As cystoscopy results were positive, a transurethral resection of the bladder tumor (TURBT) was performed. Histopathological analysis documented a Stage T1 BC. The patient’s urine sample was collected prior to medical intervention (I1), and the urine proteome was analyzed (see Fig. 4A for the comprehensive timeline). On follow-up, the patient had three more TURBT procedures within four months as symptoms worsened. A second urine sampling was taken after the fourth TURBT (I2). The two urine proteomes were compared and the dPPi calculated (Equation 1). The dPPi at the time of recurrent disease (I2) was 47% higher than the dPPi at the onset of disease (Fig. 4E, I1 - I2). The proteomes of both urines were similar with little difference in the numbers of proteins differentially expressed (Fig. 4B, I1 *vs* I2). However, the levels of the proteins were higher (Fig. 4F, PI). Remarkably, the eight pathways selected to follow the development of the disease were all overexpressed five months after the first intervention (Fig. 4E, I1 - I2), thus confirming the hypothesis that the urine proteome reflects the changes in the patient’s health after BC onset. The pertinence of the dPPi approach is illustrated by the variation observed in the MAPK signaling pathway (P8 in Fig. 4E, and 4F).

**Figure 4.**
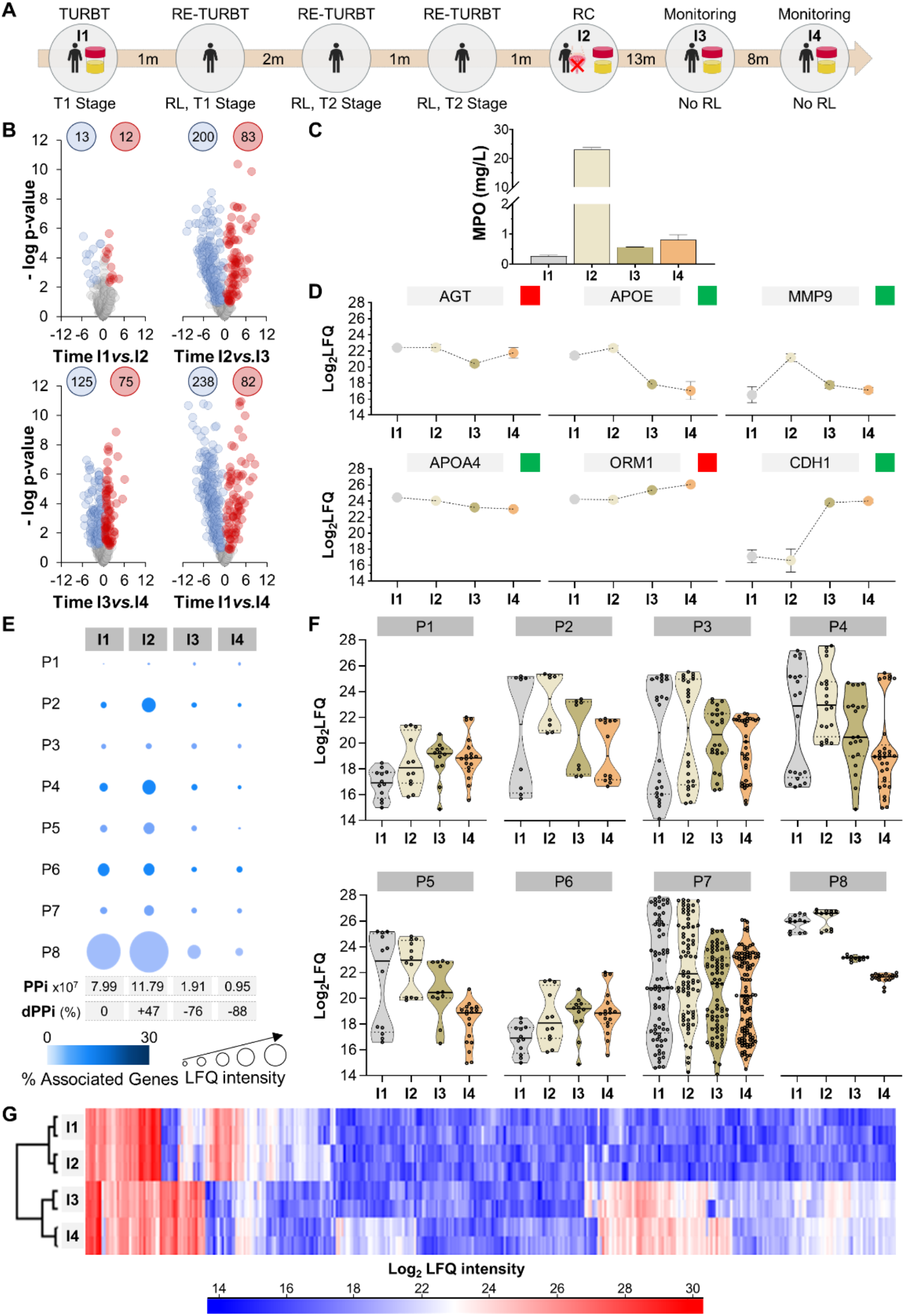
**a**, Timeline of disease course in Patient III showing intervals in months (m) between medical interventions and urine sampling (I1to I4). TURBT: trans urethral resection of bladder tumor; Re-TURBT: repeated trans urethral resection of bladder tumor; RL: relapse; RC: radical cystectomy. **b**, Volcano plots showing proteome changes in the urines sampled at times I1, I2, I3 and I4. Dots represent proteins showing statistically significant increases (red), decreases (blue) or non-statistically significant changes (gray) at the different timepoints according to Student’s *t*-test (FDR 0.05 and S0 of 0.1). **c**, ELISA quantification of Myeloperoxidase (MPO, ng/mL) in urine samples. **d**, Variation of six known protein biomarkers for bladder cancer: Angiotensinogen (AGT)^7^, Apolipoprotein E (APOE)^8^, Matrix metalloproteinase-9 (MMP9)^9^, Apolipoprotein A-IV (APOA4)^10^, Alpha-1-acid glycoprotein 1 (ORM1)^11^ and Cadherin-1 (CDH1)^12,13^. Green and red squares indicate whether the variation in the urine of Patient I accords or does not accord, respectively, with trends reported for each marker in the literature. **e**, Personal pathway index (PPi) for each time point and estimated differential PPi (dPPi) calculated as explained in the text. P1: Interleukin-12-mediated signaling pathway; P2: endodermal cell differentiation; P3, proteoglycan binding; P4: hydrogen peroxide catabolic process; P5: peroxidase activity; P6: Complement cascade; P7: humoral immune response; P8: Oncogenic MAPK signaling. **f**, Variation in protein LFQ values (including replicates) at each sampling time point for each pathway assessed. **g**, Hierarchical clustering of the four urinary proteomes of Patient I. Protein LFQ values were used to perform the cluster analysis (with average linkage, no constraint, preprocessing with k-means and Euclidean distance between column trees).

This pathway has been reported to be involved in BC development^14^and overexpression of MAPK signaling proteins has been linked to mutation in the P53 gene and to poor prognosis^15^. At this stage (I2) the medical decision was radical cystectomy. The urine proteome was analyzed again eight months and eleven months after surgery, (Fig. 4E, I3) and (Fig. 4E, I4) respectively. The comparison of the four proteomes since the onset of disease (Fig. 4G) clearly shows clustering of the urines into two groups, one before and other after radical cystectomy. This reflects the temporal differences in the pathways’ signature between both urine groups based on personal response to disease evolution and medical care.

Interestingly, the expression levels of all the pathways decreases after resection of the bladder (I3 and I4 in Fig. 4E and Fig. 4F). The dPPi decreased 76% between onset and I3, when urine was not in contact with the bladder, and then decreased 50% between I3 and I4 (Fig. 4E), the latter being consistent with the recovery experienced by this patient. Currently, this patient is in good health.

Here we have demonstrated how assessment of the urine proteome using the dPPi algorithm is a robust tool to monitor the course of BC disease and guide clinical decision-making. As the disease progresses, the levels of the proteins in hallmark pathways change. Perturbations in certain pathways at different stages of disease are described in the literature, and this information guided the choice of which pathways to include in formulating the dPPi algorithm. Equally, a preliminary study comparing different stages of the disease could be done to uncover the pathways and proteins affected. The dPPi concept can be extended to other diseases and other biopsies, like serum, plasma, or saliva, to obtain a useful readout reflecting changes in the proteome due to the influence of the disease process as well as of medical care.

## Methods

### Patients

Human mid-stream urine specimens were collected from patients whose data is reported in Table SM 1. In all cases bladder cancer diagnosis was confirmed by pathological examination. All patients were informed about the study and provided a signed informed consent according with the Central Lisbon Hospital Center Ethics Committee. Selection of patients to enroll the study was based on the following criteria: (a) Inclusion: clear bladder cancer diagnosis, and (b) Exclusion: no records of urinary cancer history, no HIV, no organ transplant, and no recent chemo/radiotherapy.

### Sample treatment

Urine samples were collected in centrifuge tubes of 50 mL (DB Falcon) containing 38 mg of boric acid (Sigma-Aldrich) in order to prevent bacterial growth^16^. Urine samples were centrifuged at 5000 × g for 20 min to remove cell debris and 10 mL aliquots of the resulting supernatants were stored at −60 °C until further use. An aliquot of urine (10mL) was concentrated by centrifugal ultrafiltration using a VivaSpin 15R (10kDa MWCO, Sartorious) at 5000g for 20 min. Finally, the concentrated urinary proteomes were quantified via a 96 well plate Bradford protein assay (Sigma-Aldrich) using and Bovine Serum Albumin to generate the calibration curve.

### Filter-aided sample preparation (FASP)

Filter Aided Sample Preparation (FASP) was used for urinary proteome digestion, following a previous modified FASP method^1,17–19^. Briefly, protein amounts ranging from 50 and 100 μg of protein were diluted with H2O to perform 200 μL and an additional 200 μL of 8 M urea (Sigma-Aldrich), 75 mM Tris-HCl (Sigma-Aldrich), 100 mM NaCl (Sigma-Aldrich), 0.02% SDS (Sigma-Aldrich). Then this protein solution was loaded onto a VivaSpin 500 (10kDa MWCO, Sartorious) and centrifuged for 20 min at 14000 × g. Each sample were processed in duplicate. The proteins present in the membrane were washed with 200 μL of 8 M urea 25 mM AmBic solution and then centrifuged for 20 min at 14,000 × g. Protein disulfide bonds were reduced by addition of 200 μL of 50 mM dithiothreitol (DTT) in 8 M urea and 25 mM AmBic and incubated for 60 minutes at 37 °C. Then, a centrifugation of 20 minutes at 14,000 × g was done, and the sample was alkylated during 45 min in the dark by the addition of 100 μL 50 mM iodoacetamide (IAA) in 8 M urea and 25 mM AmBic solution. Subsequently, a centrifugation of 20 minutes at 14,000 × g was processed and then the samples were washed twice with 200 μL 25 mM AmBic. Finally, protein digestion was performed by addition of 100 μL of trypsin solution (1:30 trypsin:protein ratio) prepared in 12.5 mM AmBic. Protein digestion was proceeded overnight (approximately 16 hours) at 37 °C and then the peptides were collected by 20 min centrifugation at 14,000 × g following by two additional membrane washing steps with 100 μL of 3% (v/v) acetonitrile containing 0.1% (v/v) aqueous formic acid and centrifugation of 20 minutes at 14,000 × g. Finally, peptides were transferred to a 500 μL microtube, dried and stored at −20 °C until further analysis by Nano-LC-MS/MS.

### LC-MS/MS

LC-MS/MS analysis was carried out using an Ultimate 3000 nano LC system coupled to an Impact HD (Bruker Daltonics) with a CaptiveSpray nanoBooster using acetonitrile as dopant. Peptides were resuspended in 100 μL of 3% (v/v) acetonitrile containing 0.1% (v/v) aqueous formic acid (FA). Protein digests were resuspended in 100 μL of 3% (v/v) acetonitrile containing 0.1% (v/v) aqueous formic acid (FA) and sonicated for 10 min using an ultrasonic bath at 100% ultrasonic amplitude, 35 kHz ultrasonic frequency. Afterwards, samples were quantified using a pierce™ quantitative colorimetric peptide assay. Afterwards, 3 μL of protein digest, containing 563 ng of peptides were loaded onto a trap column (Acclaim PepMap100, 5 μm, 100 Å, 300 μm i.d. × 5mm) and desalted for 5 min from 3% to 5% B (B: 90% acetonitrile 0.08% FA) at a flow rate of 15 μL.min^−1^. Then the peptides were separated using an analytical column (Acclaim™ PepMap™ 100C18, 2 μm, 0.075 mm i.d × 150 mm) with a linear gradient at 300 nL.min^−1^(mobile phase A: aqueous FA 0.1% (v/v); mobile phase B 90% (v/v) acetonitrile and 0.08% (v/v) FA) 5-90 min from 5% to 35% of mobile phase B, 90-100min linear gradient from 35% to 95% of mobile phase B, 100-110 95% B. Chromatographic separation was carried out at 35 °C. MS acquisition was set to cycles of MS (2 Hz), followed by MS/MS (8-32 Hz), cycle time 3.0 s, with active exclusion (precursors were excluded from precursor selection for 0.5 min after acquisition of 1 MS/MS spectrum, intensity threshold for fragmentation of 2500 counts). Together with active exclusion set to 1, reconsider precursor if the intensity of a precursor increases by a factor of 3, this mass was taken from temporarily exclusion list and fragmented again, ensuring that fragment spectra were taken near to the peak maximum. All spectra were acquired in the range 150-2200 m/z. The mass spectrometry proteomics data have been deposited to the ProteomeXchange Consortium^20^via the PRIDE^21^partner repository with the data set identifier PXD025139. To validate Myeloperoxidase label-free quantification levels, a validation was performed, using a commercially available ELISA kit (ab136943 - Myeloperoxidase (MPO) Human ELISA kit from abcam). The method was carried out according to the standard protocol described by the manufacture.

### Bioinformatics

Relative label-free quantification was carried out using MaxQuant software V1.6.17.0. All raw files were processed in a single run with default parameters^22,23^. Database searches were performed using Andromeda search engine with the were searched against the Swiss-Prot database 57.15 (515,203 sequences; 181,334,896 residues), setting the taxonomy to Human (20,266 sequences)^24^. Data processing was performed using Perseus (version 10.6.10.5) with using default settings using the workflow depicted in Fig. 1B^25^. Protein group LFQ intensities were log2-transformed, and the quantitative profiles were filtered for missing values with the following settings: min valid percentage of 50% in at least one group and values greater than 0. To overcome the obstacle of missing LFQ values, missing values were imputed using the parameters, with = 0.5 and down shift = 1.8. Log ratios were calculated as the difference in average log2 LFQ intensity values between the tested conditions (two-tailed, Student’s *t*-test, permutation-based FDR 0.05 and S0 of 0.1). Perseus was also used to obtain clusters, using average linkage, no constraint, preprocess with k-means and euclidean distance between column trees.

## Supporting information

Table SM 1, clinical information about T1 BCa groups

Table SM 2, proteins that were differentially expressed between the BCa T1 groups

## Supplementary data

Table SM 1, clinical information about T1 BCa groups.

Table SM 2, proteins that were differentially expressed between the BCa T1 groups.

## Author Contributions

J.L.C., C.L., L.C.P. and H.M.S. designed the experimental work and provide financial support. L.B.C. performed the laboratorial work and data analysis under the supervision of H.M.S. J.L.C. and H.M.S drafted the manuscript. J.L.C., C.L.E., L.C.P, R.D., M.M. and H.M.S. revisited the drafted version, corrected it and made valuable suggestions. L.C.P. and M.M. managed patient interventions and provided samples and medical data.

## Acknowledgements

PROTEOMASS Scientific Society is acknowledged by the funding provided to the Laboratory for Biological Mass Spectrometry Isabel Moura. Authors acknowledge the funding provided by the Associate Laboratory for Green Chemistry LAQV which is financed by national funds from FCT/MCTES (UIDB/50006/2020 and UIDP/50006/2020). H. M. S. is funded by the FCT 2015 Investigator Program (IF/00007/2015). L. B. C. is funded by the FCT PhD grant 2019 (SFRH/BD/144222/2019). The mass spectrometry proteomics data have been deposited to the ProteomeXchange Consortium^20^via the PRIDE^21^partner repository with the data set number PXD025139.

