## Supplementary material for "Pathway centered analysis to guide clinical decision-making in precision medicine": Table SM 1, clinical information about T1 BCa groups

| Patient Name | Status | Gender | Patient Age at Urine Collection | Histology (Stage T) | Macroscopy | Complete resection | Bisected muscularis propria | Re-TURBT | Grade | Time of Recurence | Time of Progression | Global survival |
|---|---|---|---|---|---|---|---|---|---|---|---|---|
| A1 | Deceased | M | 61 | 1 | 1, >3cm | Y | Y | N | High | 6 | 16 | 26 |
| A2 | Deceased | M | 62 | 2 | - | - | - | - | - | 9 | 9 | 10 |
| B | Alive | M | 56 | 1 | 1, >3cm | Y | Y | Y | High | N | N | 12 |
| C | Alive | M | 78 | 1 | >3, <3cm | Y | Y | - | High | 5 | N | 60 |
| D | Alive | M | 74 | 1 | >3, <3cm | Y | Y | N | High | 11 | 48 | 48 |
| E | Alive | M | 77 | 1 | 1, >3cm | Y | N | - | High | 13 | N | 15 |
| F | Alive | M | 79 | 1 | >3, >3cm | Y | N | - | High | N | N | 10 |
| G | Deceased | M | 81 | 1 | - | Y | N | Y (T2) | High | 1 | 1 | 9 |
| H | Deceased | M | 91 | 1 | 1, >3cm | N | Y | N | High | 3 | 3 | 25 |
| I | Alive | M | 78 | 1 | 1, >3cm | N | Y | Y | High | 16 | 16 | 30 |
| J | Alive | M | 74 | 1 | >3, >3cm | N | Y | Y | High | 7 | 13 | 31 |
| K | Deceased | M | 75 | 1 | 1, >3cm | Y | Y | Y (No tumor) | High | N | N | 25 |
| L | Alive | M | 79 | 1 | - | - | - | - | Low | N | N | 24 |
| M | Alive | M | 47 | 1 | 2, <3cm | Y | N | Y | Low | - | - | - |
| N | Alive | M | 65 | 1 | 1, <3cm | - | - | - | High | N | N | 50 |
| O | Alive | M | 68 | 1 | 1, <3cm | Y | Y | - | High | 9 | 9 | 9 |
| P | Alive | M | 74 | 1 | >3, >3cm | - | - | - | High | N | N | 44 |

| Drug | K | L | M | N | O | P | A1 | A2 | B | C | D | E | F | G | H | I | J |
|---|---|---|---|---|---|---|---|---|---|---|---|---|---|---|---|---|---|
| BCG Treatment | | | ■ | | ■ | | ■ | | | | ■ | ■ | | | | | |
| simvastatin | | | | | | ■ | ■ | ■ | | | ■ | ■ | | | ■ | ■ | |
| omeprazole | | | | | | ■ | ■ | ■ | | | ■ | | | | | | ■ |
| metformin | | ■ | | | | | | | | | ■ | | | | | ■ | ■ |
| losartan | | | | | | ■ | | | | ■ | | | | | | | |
| brinzolamide | | | | ■ | | | | | | | ■ | | | | | | |
| tamsulosin | | | | | | | | | | | | | ■ | | | | ■ |
| furosemide | | ■ | | | | | | | | | | | | | | | ■ |
| allopurinol | | ■ | | | | | | | | | | | | | | | ■ |
| fenofibrate | | ■ | | | | | | | | | | | | | | | ■ |
| amlodipine | ■ | | | | | ■ | | | | | | | ■ | | | | |
| bisoprolol | ■ | | | | ■ | | | | | | | | | | | | |
| acetylsalicylic acid | | ■ | | | | ■ | | | | | | | | | | | |
| ibersatan | | | | | | | | | | | ■ | | | | | | |
| silodosin | | | | | | | | | | | ■ | | | | | | |
| permixon | | | | | | | | | | | ■ | | | | | | |
| naphthidrofuril | | | | | | | | | | | | | ■ | | | | |
| perindopril | | | | | | | | | | | | | ■ | | | | |
| timolol | | | | | | | | | | | | | ■ | | | | |
| travoprost | | | | | | | | | | | | | ■ | | | | |
| indacaterol | | | | | | | | | | | | | | ■ | | | |
| budesonide | | | | | | | | | | | | | | ■ | | | |
| lercanidipine | | | | | | | | | | | | | | | ■ | | |
| irbesartan | | ■ | | | | | | | | | | | | | ■ | | |
| alprazolam | | | | | | | | | | | | | | | ■ | | |
| finasteride | | | | | | ■ | | | | | | | | | | | ■ |
| atorvastatin | | | | | | | | | | | | | | | | | ■ |
| telmisartan | | | | | | | | | | | | | | | | | ■ |
| pregabalin | | | | | | | | | | | | | | | | | ■ |
| valdoxan | | | | | | | | | | | | | | | | | ■ |
| pantoprazole | ■ | | | | | | | | | | | | | | | | |
| levitiracetam | ■ | | | | | | | | | | | | | | | | |
| dabigratano | ■ | | | | | | | | | | | | | | | | |
| pitavastatin | ■ | | | | | | | | | | | | | | | | |
| quetiapine | ■ | | | | | | | | | | | | | | | | |
| perindonpril | ■ | | | | | | | | | | | | | | | | |
| diltiazem | | ■ | | | | | | | | | | | | | | | |
| hydrochlorothiazide | | ■ | | | | | | | | | | | | | | | |
| formoterol | | ■ | | | | | | | | | | | | | | | |
| glucosamine | | ■ | | | | | | | | | | | | | | | |
| tiotropium bromide | | ■ | | | | | | | | | | | | | | | |
| acetazolamide | | | | ■ | | | | | | | | | | | | | |
| peridonpril | | | | | ■ | | | | | | | | | | | | |
| indapamide | | | | | ■ | | | | | | | | | | | | |

■ Positive

□ Negative
